# RNA-binding proteins and regulatory networks involved in life-stage, stress temperature, and drug resistance in *Leishmania parasites*

**DOI:** 10.64898/2026.02.16.706094

**Authors:** J. Eduardo Martinez-Hernandez, Victor Aliaga Tobar, Almendra Hidalgo-Cabrera, Jose M. Requena, Rubens Monte-Neto, Vinicius Maracaja-Coutinho, Alberto J. M. Martin

**Affiliations:** Núcleo de Investigación en Data Science, Facultad de Ingeniería y Negocios, Universidad de Las Américas, Santiago 7500975, Chile; Centro de Genómica y Bioinformática, Facultad de Ciencias, Ingeniería y Tecnología, Universidad Mayor, Santiago 8580745, Chile; Centro de Biologia Molecular Severo Ochoa (CBM), CSIC - Universidad Autonoma de Madrid, c/ Nicolás Cabrera, 1, 20849 Madrid, Spain; Biotecnologia Aplicada ao Estudo de Patógenos, Instituto René Rachou, Fundação Oswaldo Cruz - Fiocruz, Belo Horizonte, Minas Gerais, Brazil; Unidad de Genómica Avanzada (UGA), Facultad de Ciencias Químicas y Farmacéuticas, Universidad de Chile, Santiago, Chile; Laboratório de Medicina e Saúde Pública de Precisão (MESP2), Instituto Gonçalo Moniz, Fundação Oswaldo Cruz, Fiocruz, Salvador, Brazil; Instituto Nacional de Ciência e Tecnologia em Saúde Digital (INCT-DigiSaúde), Salvador, Brazil; Laboratorio de Redes Biológicas, Fundación Ciencia & Vida, Santiago, Chile; Escuela de Ingeniería, Facultad de Ingeniería, Universidad San Sebastián, Santiago, Chile

**Keywords:** Leishmania, gene expression regulation, post-transcriptional regulation, transcription factors, RNA binding proteins, epitranscriptomics, antimony resistance

## Abstract

Trypanosomatid parasites of the genus *Leishmania* rely on post-transcriptional regulation of gene expression because gene transcription is not canonically regulated at the single-gene level. RNA-binding proteins (RBPs) are central to this regulatory architecture, yet their genome-wide diversity, conservation, and condition-specific associations remain incompletely defined across the genus. Here, we combined comparative genomics and systems-level transcriptomic analyses to map the RBP repertoire in 33 strains spanning 19 *Leishmania* species. We also connect RBPs to developmental, stress, and drug-resistance contexts, using Leishmania braziliensis as an example. Using probabilistic domain detection and homology-based annotation, we identified 38,662 putative RBPs across all genomes, with 1,114 to 1,491 RBPs per genome. Comparative genomics analysis revealed a conserved core of 404 RBP clusters shared across all examined species, alongside lineage-restricted clusters in major *Leishmania* groups. We notably detected widespread conservation of enzymes linked to messenger RNA modification (for example, NAT10, TRMT6/61A, and NSUN2), but failed to identify canonical N6-methyladenosine writer orthologs, suggesting divergence of this machinery across *Leishmania* genomes. In *L. braziliensis*, expression profiling in different life-cycle stages and stress conditions highlighted stage-biased RBPs, including elevated ZC3H20 in amastigotes and increased RBP6 in metacyclic promastigotes. Finally, co-expression network analysis of trivalent antimony (SbIII) resistant genotypes identified RBPs co-expressed with genes previously associated with antimony resistance. In contrast, motif-based analysis supported a putative DRBD3-centered post-transcriptional module that includes 10 candidate stabilized transcripts in SbIII-resistant promastigotes. Together, these results provide a comparative framework to prioritize RBPs and associated regulatory programs implicated in parasite adaptation and antimony resistance.

**AUTHOR SUMMARY:** Leishmaniasis results from infection by Leishmania parasites and remains challenging to control due to limited treatments and the continual emergence of drug resistance, especially to antimonials. These parasites exhibit an unusual gene regulation method; instead of activating or deactivating transcription for individual genes, they depend heavily on proteins that bind RNA to determine which messages are retained, translated, or degraded. In this study, we created a comprehensive atlas of RNA-binding proteins across the Leishmania genus by analyzing 33 parasite strains from 19 species. We identified a large core set of conserved RNA-binding proteins shared by all species, along with lineage-specific proteins that may help different parasite groups adapt to various hosts and environments. Additionally, we mapped enzymes responsible for RNA chemical modifications and found that the typical machinery for m6A marks in many organisms appears to be divergent in *Leishmania*. Finally, by associating RNA-binding proteins with specific life-cycle stages, stress responses, and antimony resistance in *L. braziliensis*, we pinpointed candidate regulators and gene modules for further experimental validation. This resource helps prioritize regulatory factors that could drive parasite adaptation and resistance.

## INTRODUCTION

*Leishmania* spp. is a genus of trypanosomatids responsible for a group of diseases collectively known as leishmaniasis (1, 2). These diseases are classified according to their wide clinical manifestations, ranging from self-healing skin ulceration (cutaneous leishmaniasis or CL) to the potentially fatal visceral form (visceral leishmaniasis or VL), which affects organs such as the liver, spleen, and bone marrow (1, 3). Current therapies are limited, often fail due to the emergence of resistant strains, and the development of new drugs remains scarce (4–6).

As early-diverging eukaryotes (7, 8), trypanosomatids are characterised by an unusual gene regulatory architecture: unlike most eukaryotes, they lack conventional transcriptional control of individual genes. Instead, gene expression is largely governed by post-transcriptional processes, including trans-splicing, polyadenylation, and stabilization of the 3′ untranslated regions (UTRs) (9, 10). Understanding these regulatory layers is crucial for determining how these parasites coordinate cellular processes that facilitate their survival, adaptation and drug resistance. In this context, low-cost computational approaches can generate insights to guide targeted experimental research.

Gene expression in *Leishmania*, as in other trypanosomatids, is primarily regulated by post-transcriptional events that result in the regulation of mRNA stability (11). Furthermore, non-coding RNAs have recently been described as potential regulators of gene expression in *Leishmania braziliensis* and *L. donovani* across their life cycle and under different stressors (12). Accordingly, gene regulation is influenced by lifestyle and the diverse stresses to which it is exposed. *Leishmania* has a digenetic life cycle, divided into vector (sandfly) and vertebrate host stages (13–15), and each environment imposes specific stressors, including changes in pH, temperature, and nutrient availability, which drive morphological transformations, biochemical adaptations, and major shifts in gene expression (16–19). For instance, nutrient stress, such as iron or other essential nutrients, requires an accurate change in gene expression to respond to this stressor (17, 20, 21). In contrast, non-coding RNA have been associated with starvation responses, parasite survival and histone modification processes (12) Among the diverse gene regulatory strategies required to survive under stress conditions, *Leishmania* and other trypanosomatids, employ post-transcriptional regulation mechanisms, including the stabilization and degradation of mRNAs mediated by RNA-binding proteins (RBPs) (22–24).

RBPs are central players in post-transcriptional regulation, compensating for the absence of canonical transcriptional control in *Leishmania* and other trypanosomatids. They act mainly through interactions with the 3′ UTRs of their target mRNAs (11, 22, 25–28). Several RBPs have been implicated in parasite differentiation. For instance, RBP10, HNRNPH, ZC3H28, and ZC3H32, are stage-specific regulators in the bloodstream form of *Trypanosoma brucei* (29–33). Others, such as ZC3H39 or ZC3H11, are essential for responses to nutrient or temperature stress (23, 34). Beyond protein-RNA interactions, chemical modifications of RNA have recently emerged as an additional layer of post-transcriptional regulation in trypanosomatids (35). Notably, RNA N6-methyladenosine (m6A) has been detected in *T. brucei*, *T. congolense*, *T. cruzi,* and *L. infantum* (36, 37). However, the enzymes responsible for m6A deposition in trypanosomatids remain unidentified, highlighting an unexplored dimension of the epitranscriptome in these organisms.

In this work, we aimed to better understand how *Leishmania* regulates gene expression through RBPs. We combined comparative genomics and systems biology approaches to identify and characterise the repertoire of RBPs across 33 strains from 19 *Leishmania* species. We further investigated their expression patterns and co-expression with protein-coding genes to uncover functional and regulatory associations with the lifecycle, temperature stress, and antimony drug resistance. Here, we present the most comprehensive comparative genomic and transcriptomic analysis of RNA-binding proteins (RBPs) and regulatory networks in *Leishmania* to date, revealing conserved regulatory elements, lineage-specific adaptations, and their involvement in stress responses and drug resistance. Surprisingly, *L. braziliensis* strains revealed multiple RBPs associated with specific lifecycle stages, and co-expressed with genes previously linked to drug resistance, underscoring the pivotal role of RBPs in the regulatory mechanisms underlying antimony resistance in *Leishmania*.

## RESULTS

### Comparative analysis of RNA-binding proteins and RNA modification machinery in *Leishmania* species

We first conducted a homology search analysis and domain characterization using probabilistic models to identify the repertoire of RBPs in *Leishmania* parasites. Through these combined search strategies, we identified a total of 38,662 putative RBPs across all evaluated *Leishmania* species. The ten most frequent motifs/domains were: 50S ribosome-binding GTPase (MMR_HSR1, 3,032), RMM domains (2,498), DEAD domains (2,250), ResIII (1,952), GTP_EFTU (1,641), RNA helicases (1,503), RIO1 family domains (1,250), methyltransferases with Methyltransf_31 domains (1,128), zinc-finger CCCH (1,123), and MTS domains (638). The total number of RBPs per genome ranged from 1,114 in *L. donovani* AG83 to 1,491 in *L. braziliensis* IOCL3564 (Supplementary File 1).

Comparative analysis revealed 404 RBP clusters that were preserved across all examined species. In contrast, 17 clusters were exclusive to the *Viannia* subgenus, another 17 to the *L. mexicana* complex, and 15 were unique to the *L. donovani* complex (Figure 1A). GO term enrichment analysis of the core RBPs (Supplementary File 2) revealed processes related to gene expression (GO:0010467), ncRNA metabolic process (GO:0034660), and ncRNA processing (GO:0034470); as well as to RNA modification activities, such as pseudouridine synthase activity (GO:0009982), methyltransferase activity (GO:0008168), and S-adenosylmethionine-dependent methyltransferase activity (GO:0008757) (Supplementary File 2). Figure 1B summarises the top 20 biological process terms enriched in the core RBPs.

**Figure 1.**
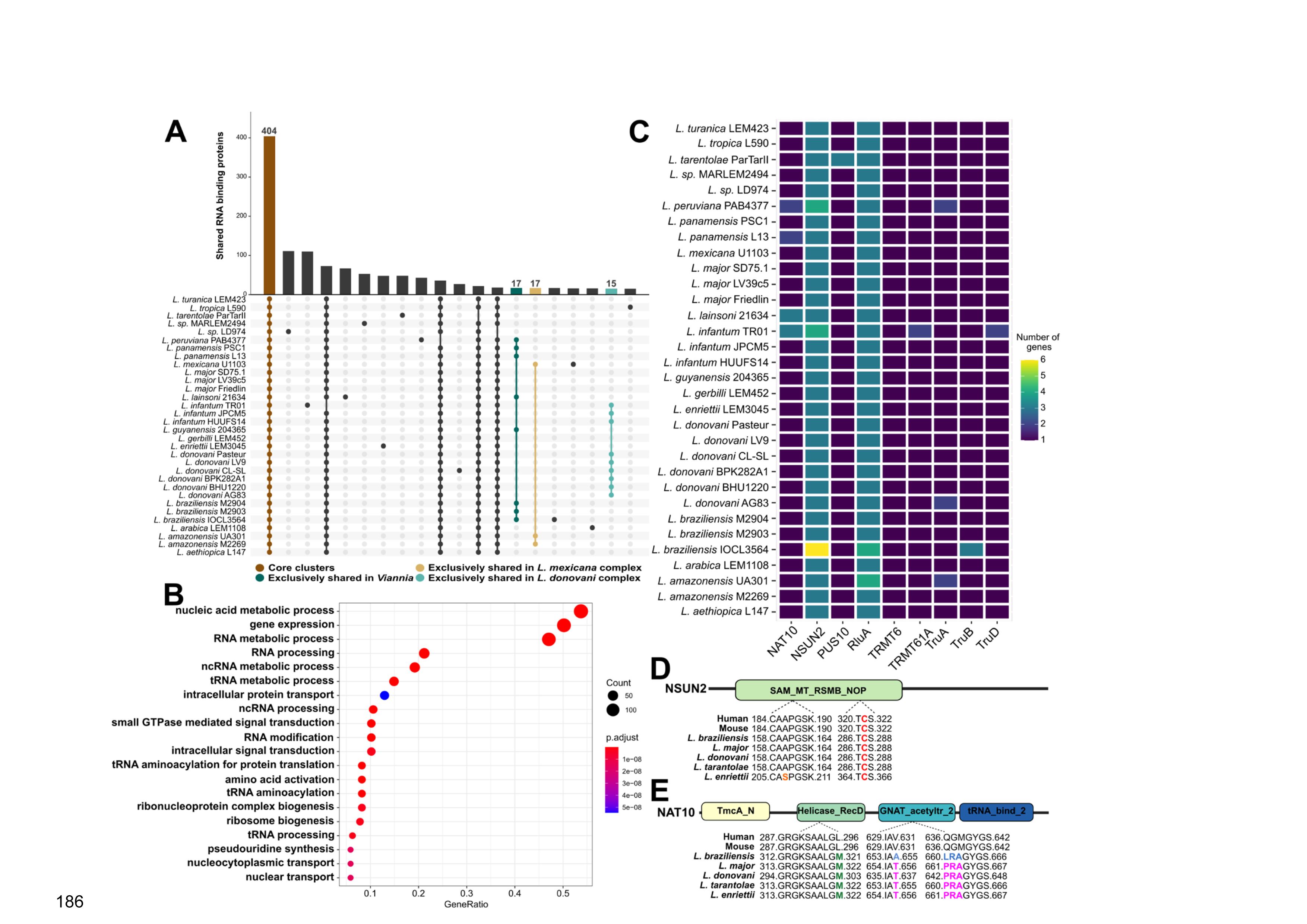
RNA-binding proteins (RBPs) across *Leishmania* species. **A**. UpSet plot showing the top 20 gene family intersections identified in the pan-RBP analysis across *Leishmania* spp. **B**. Gene Ontology (GO) enrichment analysis of biological processes associated with core RBPs conserved throughout the genus *Leishmania*. **C.** Heatmap illustrating the presence and distribution of writers RBPs identified in *Leishmania* parasites. **D.** Schematic alignment of NSUN2 orthologs comparing *Leishmania* and mammalian sequences. Conserved catalytic residues within the RsmB/F methyltransferase domain are highlighted in red. **E**. Domain architecture comparison of NAT10 orthologs in *Leishmania* spp., humans, and mice. Green indicates a Leu/Met substitution in the RNA-binding helicase domain. The blue highlights show a Val-to-Ala substitution at the third position of the mammalian motif (IAV and QGM/LRA) in the *Viannia* subgenus (*L. braziliensis*) species. In magenta, Val/Thr and QGM/PRA substitutions are shown for species within the *Leishmania* (*L. donovani* and *L. major*), *Sauroleishmania* (*L. tarentolae*), and *Mundinia* (*L. enriettii*) subgenera, located at the Acetyl-CoA binding site of the NAT10 N-acetyltransferase domain.

To explore potential links with the epitranscriptomic machinery, we aligned each predicted RBP against a reference list of RBPs focused on mRNA modification writers (a kind of mRNA binding proteins, mRBP) (35), and RBPs regulators of gene expression in trypanosomatids (25).

Thus, we identified three orthologs of the writer NOP2/SUN RNA methyltransferase family member 2 (NSUN2) in all genomes analysed, with six copies in *L. braziliensis* IOCL3564 (Figure 1C). This RBP is responsible for the methylation of m5C (5-methylcytosine), contains a methyltransferase RsmB/F domain, and was annotated as an S-adenosyl-L-methionine-dependent methyltransferase. Next, we evaluated the similarity between leishmanial NSUN2 proteins and their human and mouse orthologs. The total sequence identity ranges from 31% to 37% with their human and mouse orthologs, respectively, increasing to 41% within the RsmB domain. We also observed high conservation of the active site across all leishmanial proteins relative to their orthologs in mice and humans. Furthermore, there was a perfect conservation of the S-adenosyl-L-methionine binding site in *Leishmania* and their human and mouse orthologs, except for a Ser to Ala substitution in *L. enrietti* (see Figure 1D). Orthologs of the writers TRMT6 and TRMT61A, which form a methyltransferase complex responsible for modifying tRNAs and specific mRNAs, were also present in all evaluated species (Figure 1C). Leishmanial TRMT6 shared an average of 40% sequence identity with its human and mouse orthologs, while TRMT61A proteins had an average identity of 36% compared to their mammalian orthologs. TRMT6 contained a Gcd10p family domain at the C-terminus. We aligned this domain from all *Leishmania* genomes with the corresponding Gcd10p domains in mouse and human TRMT6 and found low conservation between mammalian and leishmanial sequences (average identity 28%). Conversely, for the GDC14 domain present in TRMT61A, we observed an average sequence identity of 41%. We further confirmed the presence of a putative ortholog of the RBP writer RNA cytidine acetyltransferase NAT10, in all *Leishmania* spp. genomes (Figure 1C and Supplementary File 1). A local alignment of leishmanial NAT10 with its human and mouse protein orthologs showed approximately 40-43% sequence identity. The identity was higher in the Helicase domain (55% sequence identity) and the DUF1726 domain (70% sequence identity), whereas the N-acetyltransferase and tRNA-binding domains exhibited lower conservation, with only 24% and 42% identity, respectively. When examining the preservation of the RNA-binding motif within the Helicase domain across leishmanial, mouse, and human NAT10 proteins, the comparison revealed perfect complementarity, with only one Leu to Met substitution in all leishmanial proteins at the last position relative to the mammalian NAT10. Conversely, we observed low conservation in the AcetylCoA binding domain, with subgenus-specific amino acid replacements: Val to Thr in species of *Leishmania*, *Sauroleishmania*, and *Mundinia* subgenera, compared to mammalian NAT10; and Val to Ala in species of the *Viannia* subgenus at the third position of motif IAV.

Additionally, some subgenus-specific substitutions were identified in the motif QGMGYGS, involved in AcetylCoA binding in humans and mice, where QGM is replaced by LRA in *Viannia* species and by PRA in the other three subgenera (Figure 1E). Subsequently, we searched for proteins that serve as Pseudouridine (Ψ) mRNA writers in the *Leishmania* species. Our results confirmed the presence of orthologs of TruA, TruB, TruD, RluA, and PUS10 in all genomes analysed. Figure 1C illustrates the total number of each RBPs identified in *Leishmania* parasites.

Finally, we conducted a sequence homology search for RBPs involved in gene expression regulation that bind mRNA (mRBPs). A total of 35 regulatory mRBPs were identified across all *Leishmania* parasites, and the functional annotation of these proteins indicated putative roles in cell cycle regulation, stage-specific regulation of procyclic and metacyclic forms, and stress response (Table 1).

**Table 1.**
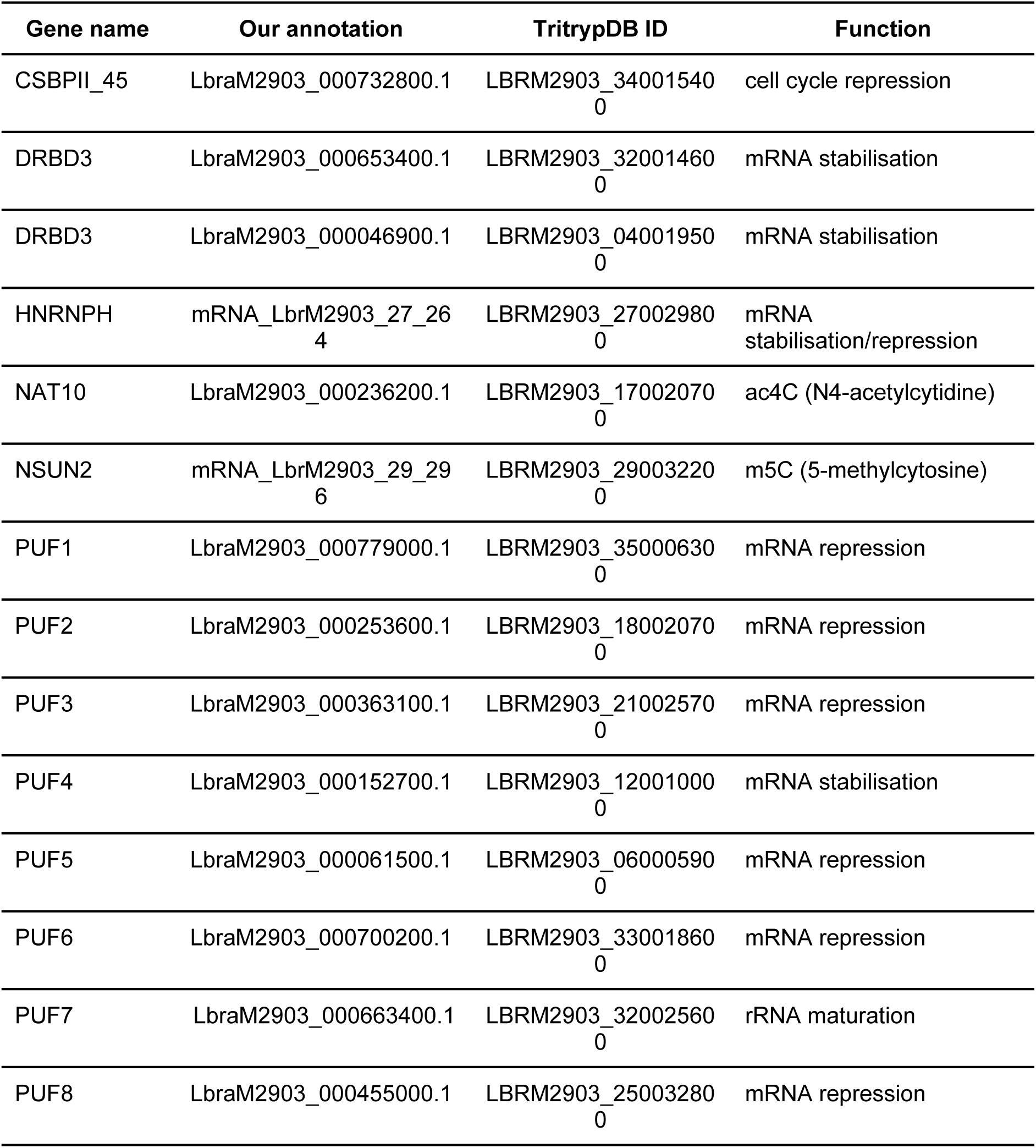
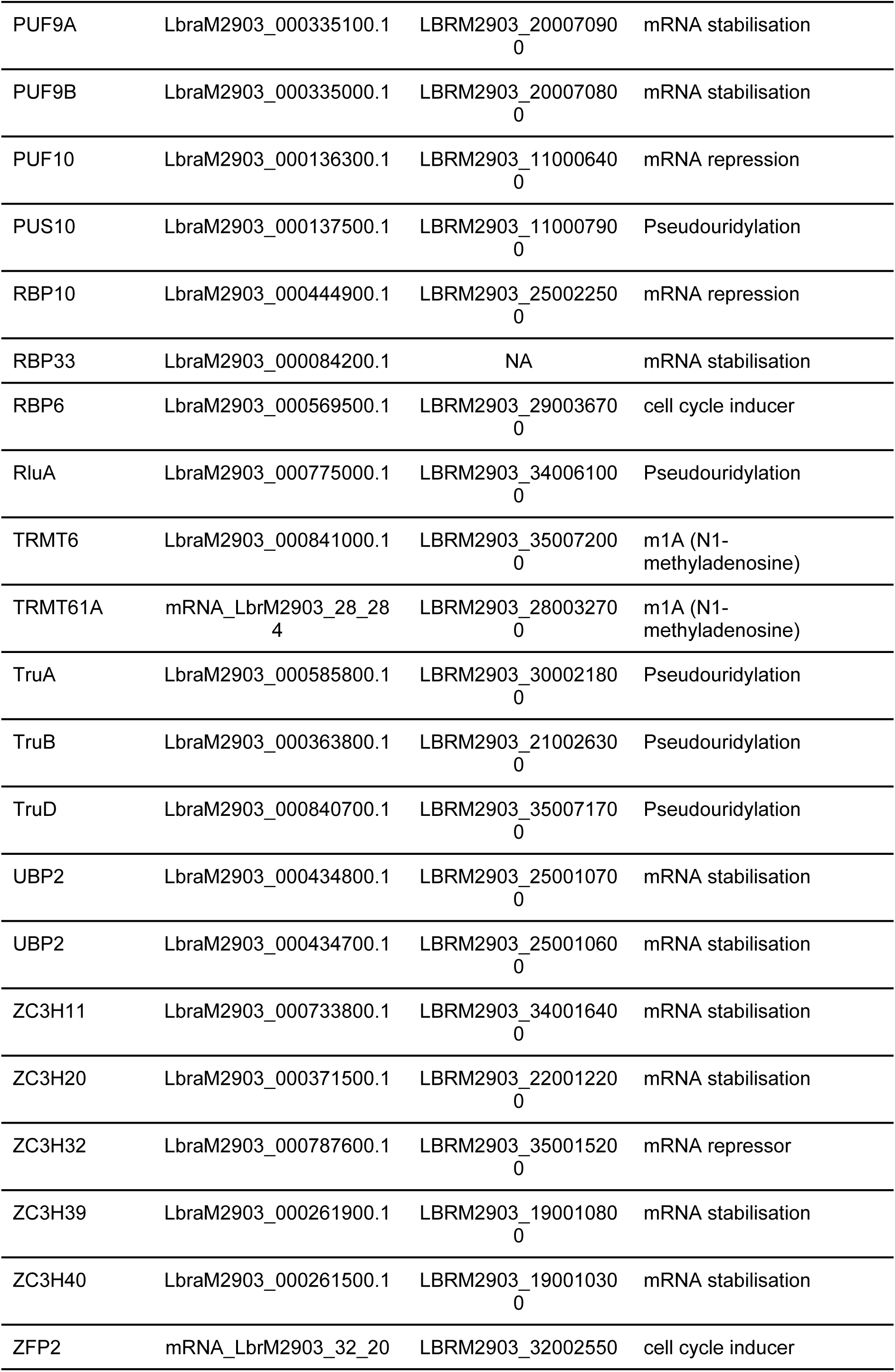
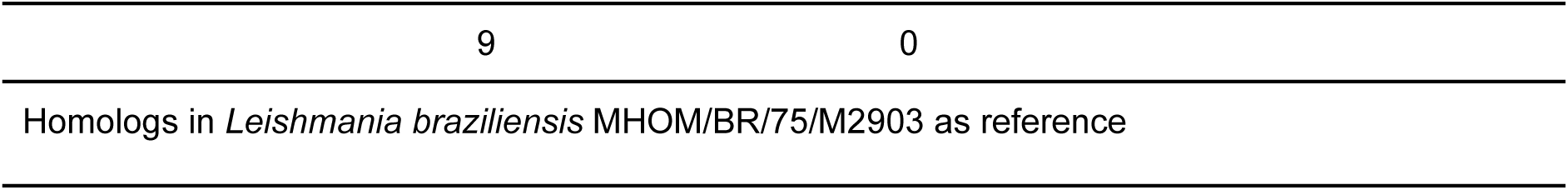
mRBPs associated with gene expression regulation and epitranscriptomics in *Leishmania* parasites.

### Differential gene expression of mRBPs across different contexts reveals stage-, stress-, and drug-specific regulation in *Leishmania braziliensis*

First, we characterised a set of differentially expressed protein-coding genes (DEGs) across three distinct contexts according to previous studies in *L. braziliensis.* These include: different developmental stages (amastigote, metacyclic and procyclic), promastigote samples under temperature stress, and promastigote samples resistance and susceptible to SbIII (antimony), to gain further insights into the potential roles of mRBPs in those processes. In total, we analysed the expression of 9,172 coding genes. Across developmental stages, we identified 1,167 genes which were exclusively overexpressed in the amastigote stage, 441 in metacyclic promastigotes, and 1,260 in procyclic promastigotes. Additionally, 848 genes were shared between the metacyclic and amastigote stages, 55 between the procyclic and amastigote stages, and 511 between the procyclic and metacyclic stages (Figure 2A, Supplementary File 3). Under temperature stress, we detected 492 genes exclusively overexpressed at 24 °C and 2,059 at 26 °C, while 180 and 538 genes were uniquely upregulated at 28 °C and 30 °C, respectively (Figure 2B, Supplementary File 3). In the context of antimony susceptibility, we identified 1,311 DEGs, including 693 upregulated in resistant promastigotes and 618 in susceptible ones exclusively (Figure 2C, Supplementary File 3). Functional enrichment analysis for each condition (Supplementary File 3) revealed distinct biological processes, with the top ten enriched GO terms illustrated in Figures 2D-F.

**Figure 2.**
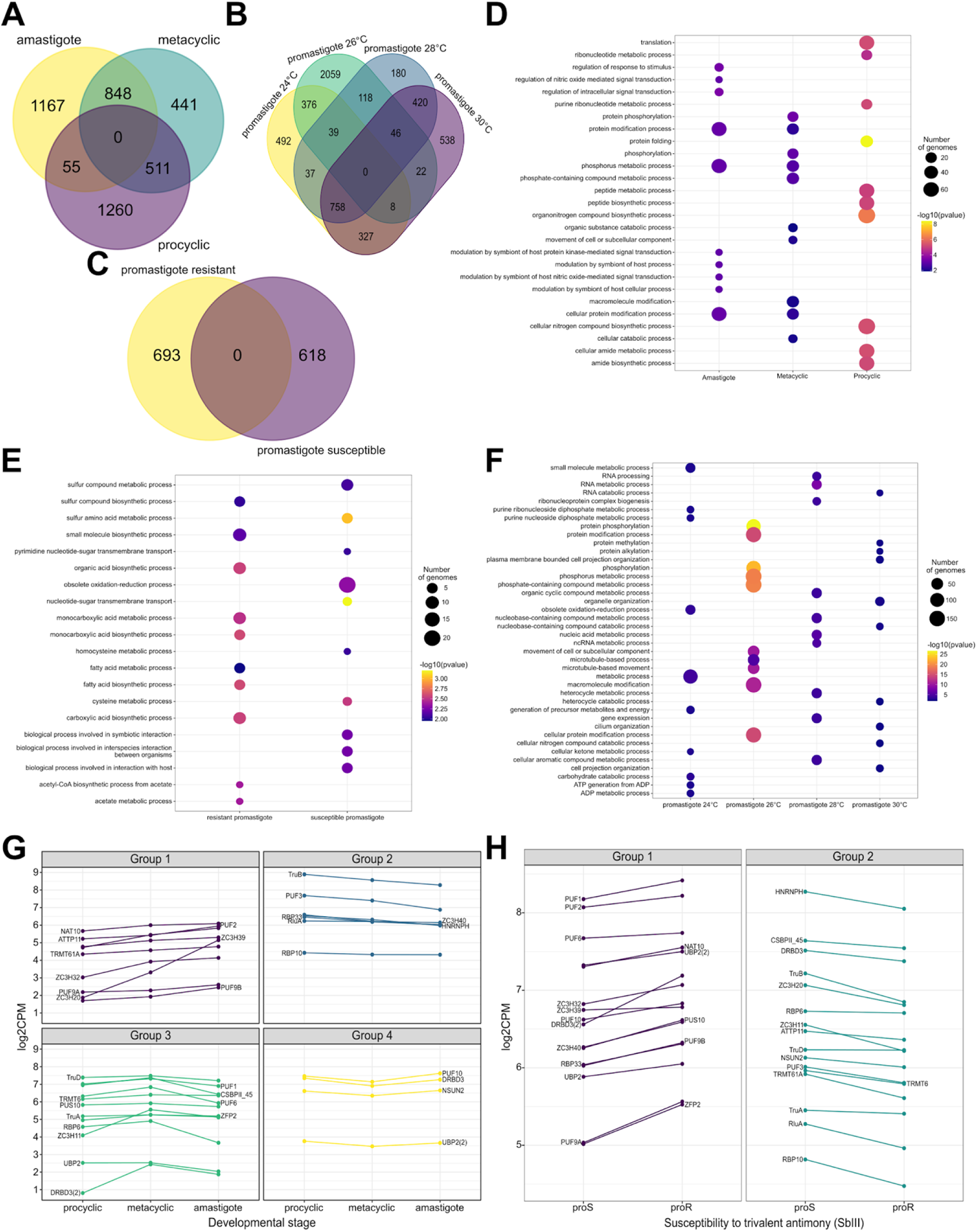
Differential Gene Expression Analysis under Various Biological and Environmental Conditions in *Leishmania braziliensis* A–C. Venn diagrams illustrating the overlap of differentially expressed genes (DEGs) in response to: **A.** developmental stage transitions (promastigote versus amastigote), **B.** temperature stress, and **C.** trivalent antimony (SbIII) resistance. **D–F.** GO enrichment analysis of biological processes linked to DEGs under each condition: **D.** developmental stage, **E.** SbIII resistance, and **F.** temperature stress. **G.** Expression profiles of selected mRNA-binding proteins (mRBPs) across the developmental stages of *L. braziliensis*. **H.** Expression profiles of selected mRBPs associated with SbIII susceptibility in *L. braziliensis* promastigotes.

Having mapped the global differential expression of protein-coding genes across developmental stages, temperature stress conditions, and SbIII susceptibility in *L. braziliensis*, we next focused on the subset of mRBPs (Table 1). Given their central role in post-transcriptional regulation, we investigated whether mRBPs exhibit specific expression patterns that could link them to life-stage regulation, temperature stress adaptation, or drug resistance. First, we identified four groups of mRBPs with distinct life stage-specific profiles. Group 1, overexpressed in amastigotes: NAT10, PUF2, PUF9, TRMT61A, ZC3H20, ZC3H32, and ZC3H39 showed elevated expression in this stage compared to metacyclic and procyclic promastigotes (Figure 2G, top left). Group 2, overexpressed in procyclic: HNRNPH, PUF3, RBP10, RBP33, RluA, TruB, and ZC3H40 were expressed at the highest levels in procyclic promastigotes, with a decrease expression in the metacyclic and amastigote stages (Figure 2G, top right). Group 3, overexpressed in the metacyclic: CSBPII_45, DRBD3, PUF1, PUF6, PUS10, RBP6, TRMT6, TruA, TruD, UBP2, and ZFP2 were highly expressed in metacyclic promastigotes (Figure 2G, bottom left). Group 4, reduce regulation in the metacyclic stage: DRBD3, PUF10, NSUN2, and UBP2 (Figure 2G, bottom right). Stage-specific expression patterns were also evident: the amastigote upregulated the ZC3H39, PUF2, ZC3H11, ZC3H20, and ZC3H32 were upregulated also in the metacyclic stage. The DRBD3 and PUF6 were uniquely overexpressed in metacyclic promastigotes, while TruB and HNRNPH were specific to procyclics. Finally, CSBPII_45, PUF3, and RBP6 were shared between procyclic and metacyclic forms. Second, under temperature stress, two major expression patterns emerged: one group peaked at 26 °C, while the other showed minimal expression at this temperature across all conditions (Supplementary Figure 1). Third, in the context of SbIII susceptibility, we identified two expression groups. The first (DRBD3, ZFP2, and PUF9A) showed strong upregulation in resistant promastigotes (Figure 4H), while the second (ZC3H11 and RBP10) displayed the most substantial expression shifts. Notably, DRBD3, ZFP2, and PUF9A were overexpressed in resistant parasites, whereas no mRBPs were specifically upregulated in susceptible ones. A complete list of the significantly expressed mRBPs for each condition is provided in Supplementary File 3.

### Context-specific co-expression networks reveal potential regulatory roles of mRBPs in *L. braziliensis*

Given the stage-, stress-, and drug-specific expression of several mRBPs, we investigated their potential regulatory roles by integrating expression patterns into context-specific co-expression networks. This approach allowed us to identify putative gene modules, regulatory hubs, and sequence motifs that may link RBPs to parasite adaptation and antimony resistance. To identify potential regulatory interactions between mRBPs and their co-expressed partners, we constructed perturbation-based co-expression networks using all 9,172 *L. braziliensis* genes that passed our counts per million (CPM) expression filter, mentioned in the materials and method section. Each context-specific network contained ∼42 million correlations. To assess network reliability, we examined 135 known amastigote biomarkers. All biomarkers were clustered within five modules (Figure 3A), and displayed preserved expression patterns throughout the lifecycle with an upregulation in amastigote stage (Figure 3B), confirming the robustness of our methodology. Of these, 122 were identified as DEGs, with 108 also differentially expressed in the metacyclic stage and 14 exclusive to amastigotes.

**Figure 3.**
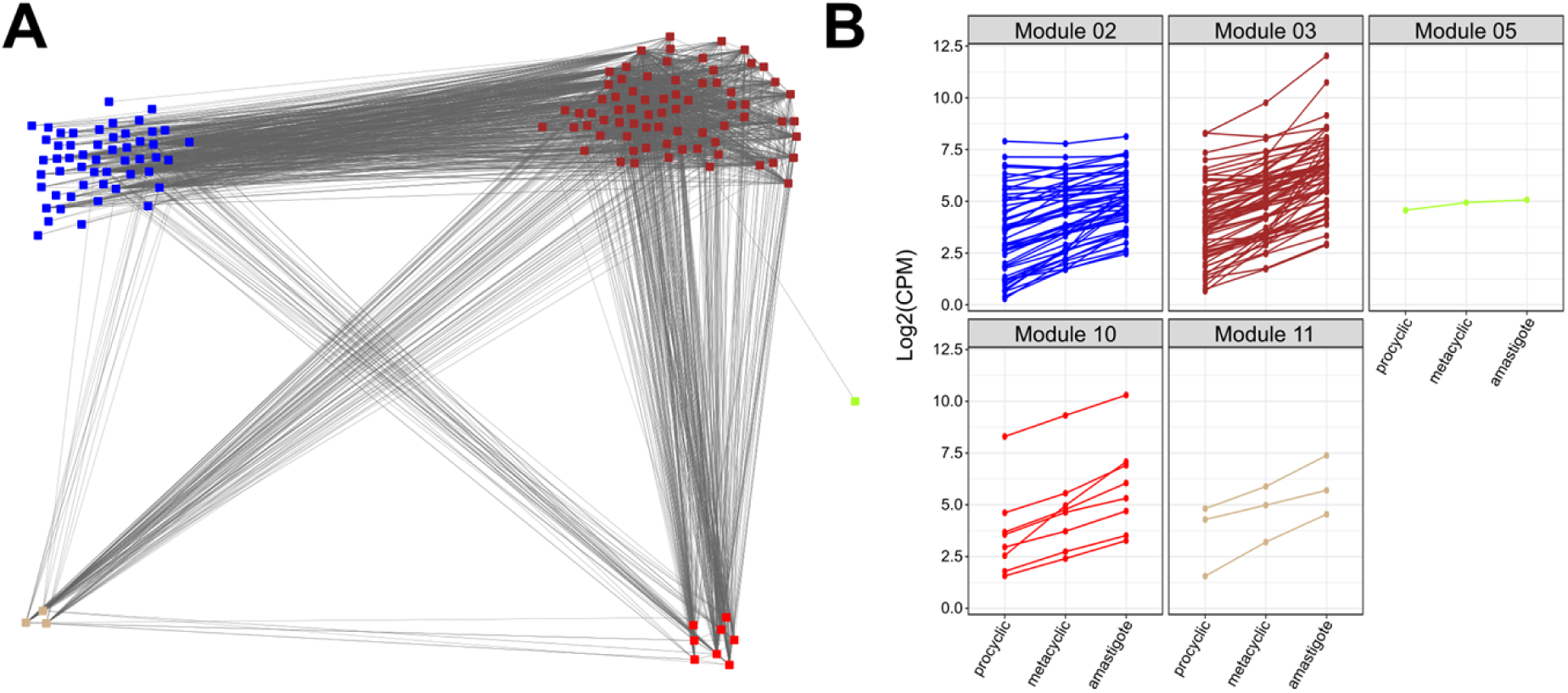
Validation of the co-expression network analysis methodology. **A.** 135 biomarkers of the amastigote form in *L. braziliensis* are categorised into the same five modules. **B.** Expression patterns of 138 amastigote biomarkers.

**Figure 4.**
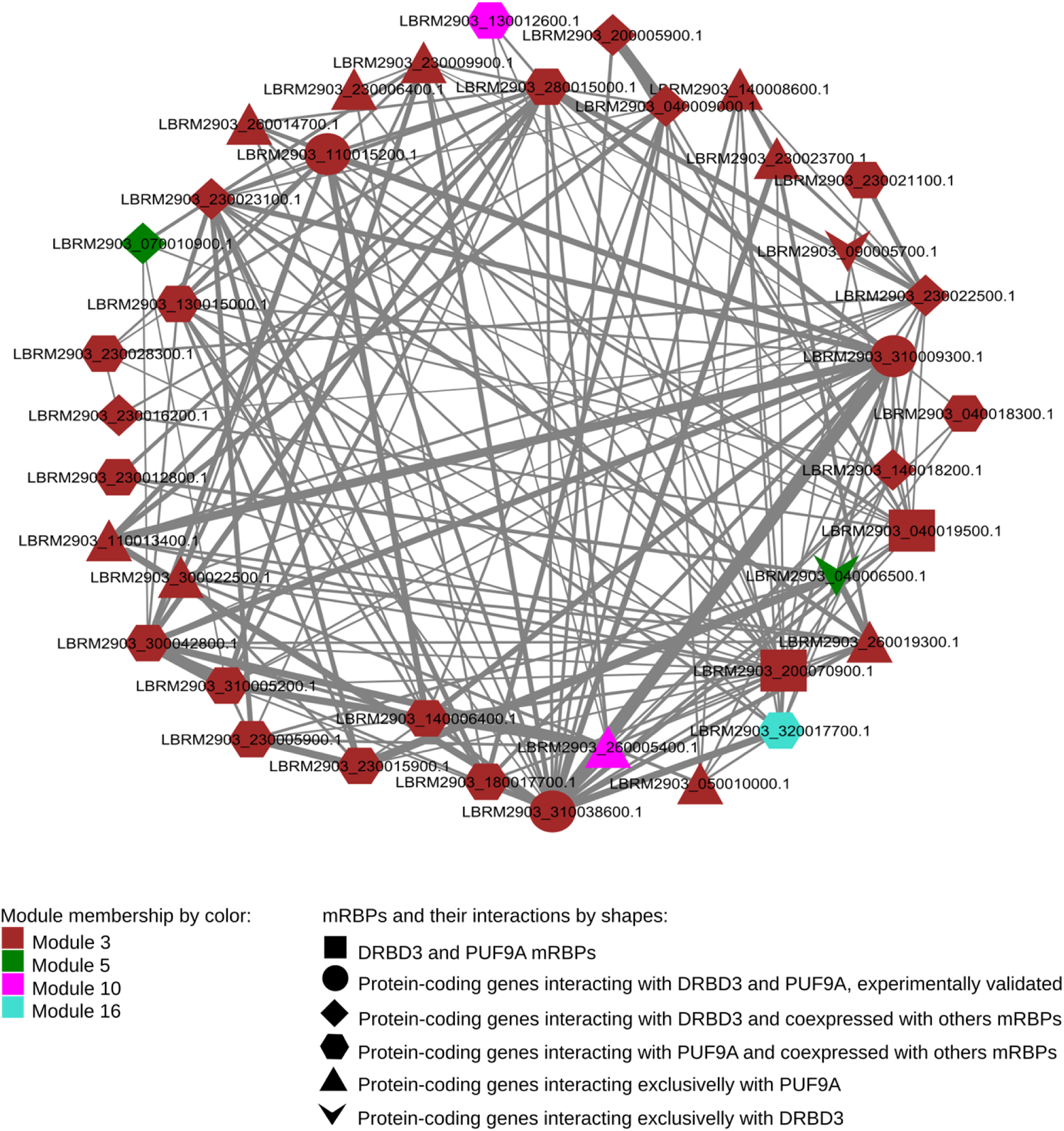
Protein-protein interaction network of co-expressed genes with DRBD3, PUF9A, and ZFP2 in SbIII-resistant *L. braziliensis* promastigotes. The colours indicate module membership and shapes represent interactions. The DRBD3 and PUF9A mRBPs are shown as rectangles, circles represent genes whose products have experimentally validated interactions with DRBD3 and PUF9A and are co-expressed with DRBD3, PUF9A, and ZFP2. Diamonds represent protein-coding genes interacting with DRBD3 and co-expressed with DRBD3, PUF9A, and ZFP2. Hexagons represent protein-coding genes interacting with PUF9A and co-expressed with DRBD3, PUF9A, and ZFP2. Triangles represent protein-coding genes interacting exclusively with PUF9A, and V-shape figures represent products co-expressed and interacting exclusively with DRBD3. Edge width corresponds to the combined score obtained from STRING DB.

As noted earlier, DRBD3, PUF9A, and ZFP2 were overexpressed in SbIII-resistant promastigotes (Figure 2H), and their products may act as stabilisers and inducers of mRNA expression (Table 1). We therefore focused on their co-expression profiles and conducted a detailed examination of how these three genes are expressed in comparison to the overexpressed coding genes in the resistant promastigotes. DRBD3 was positively co-expressed with 20 DEGs, PUF9A with 181 DEGs, and ZFP2 with 2 DEGs. Shared interactions included 114 DEGs co-expressed with both DRBD3 and PUF9A, 2 shared between PUF9A and ZFP2, and 246 DEGs co-expressed with all three mRBPs GO term enrichment for these co-expressed genes is shown in Supplementary File 4.

Subsequently, we mapped our co-expression subnetworks to the protein-protein interaction data in the STRING database to identify experimentally validated interactions between mRBPs and co-expressed coding genes. Among the 287 DEGs co-expressed with DRBD3, PUF9A, and ZFP2, three encoded proteins, one SEC61-like protein and two RNA polymerases (a DNA-directed RNA polymerase and the RNA polymerase II largest subunit, also called RBP1), were identified as experimentally validated interactors with DRBD3 and PUF9A. From the same set, we identified seven genes whose encoding proteins physically interacted exclusively with DRBD3, and 14 with PUF9A (Table 2, Figure 4). In addition, previous experimentally validated edges included two interactions between DRBD3 and proteins from its 20 co-expressed genes, and eleven interactions involving PUF9A from its 182 co-expressed DEGs (Table 2).

**Table 2.**
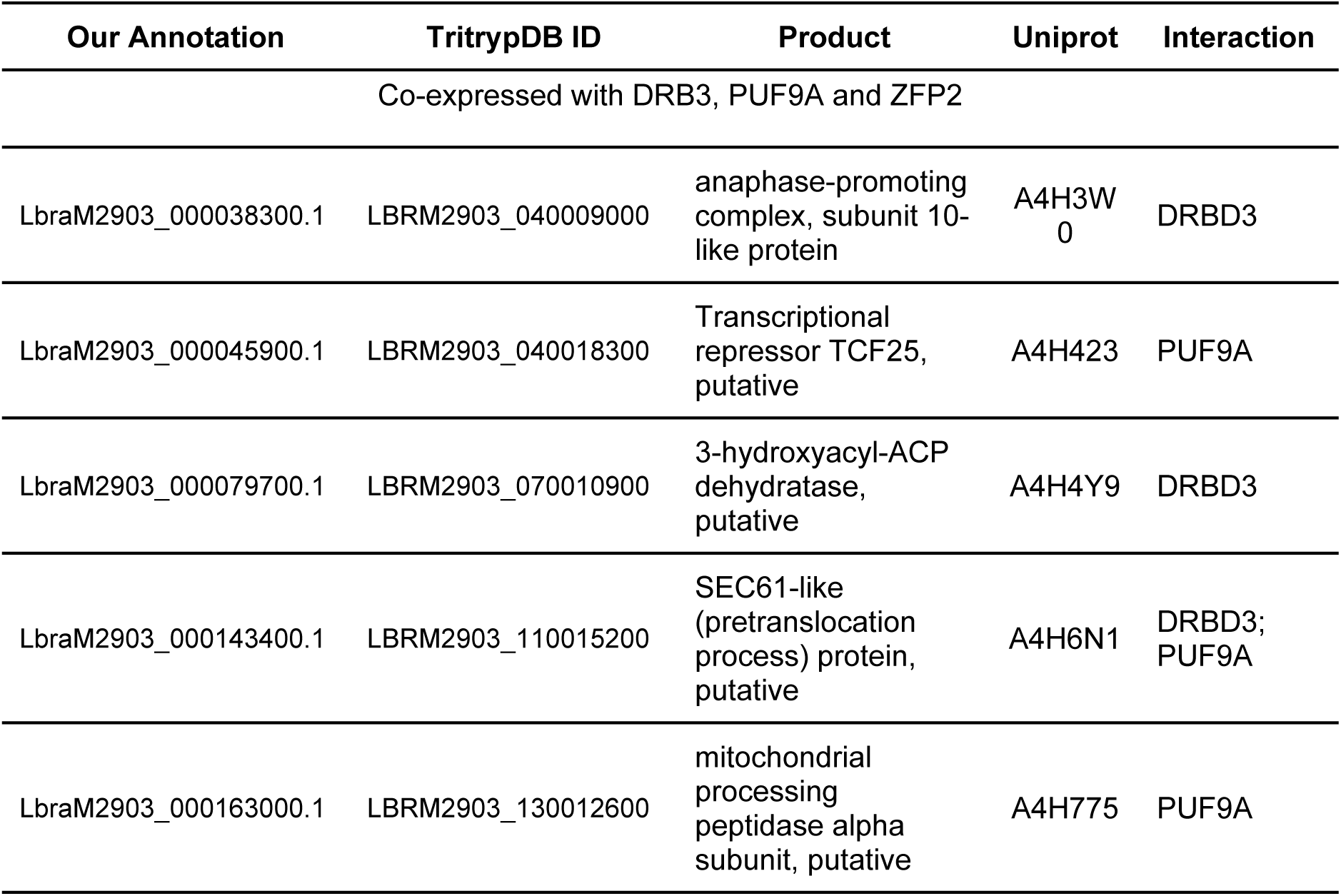
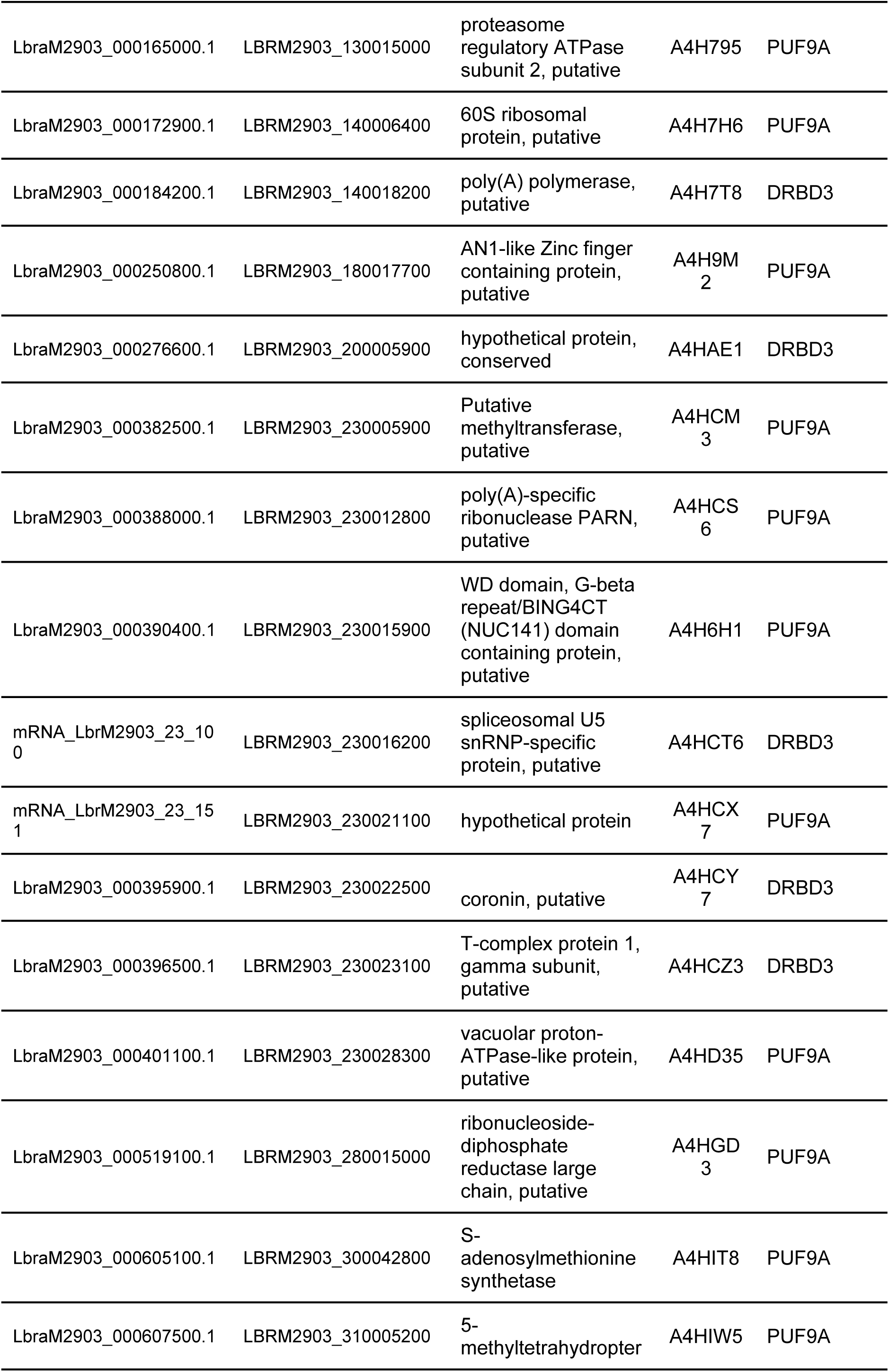
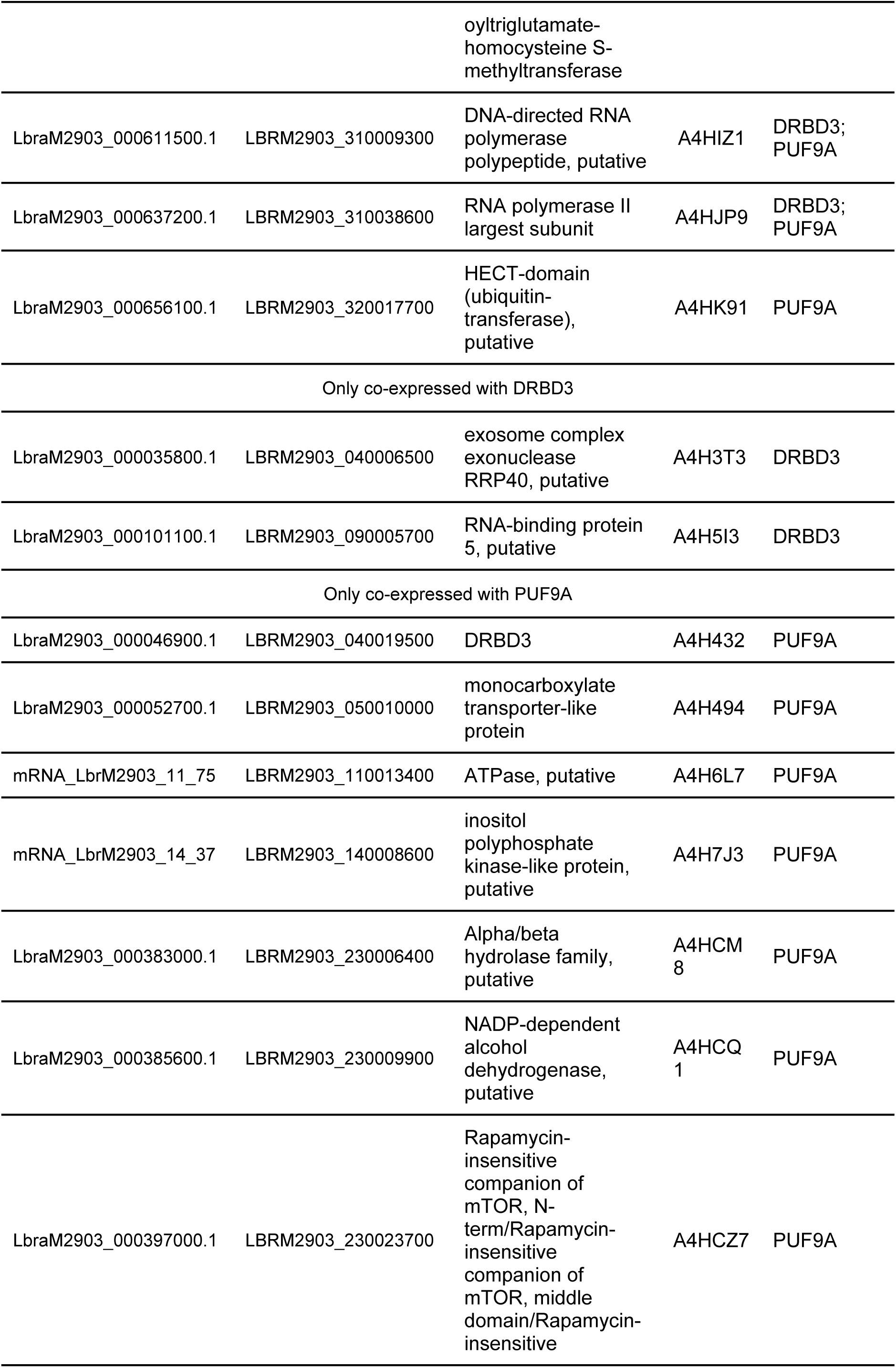
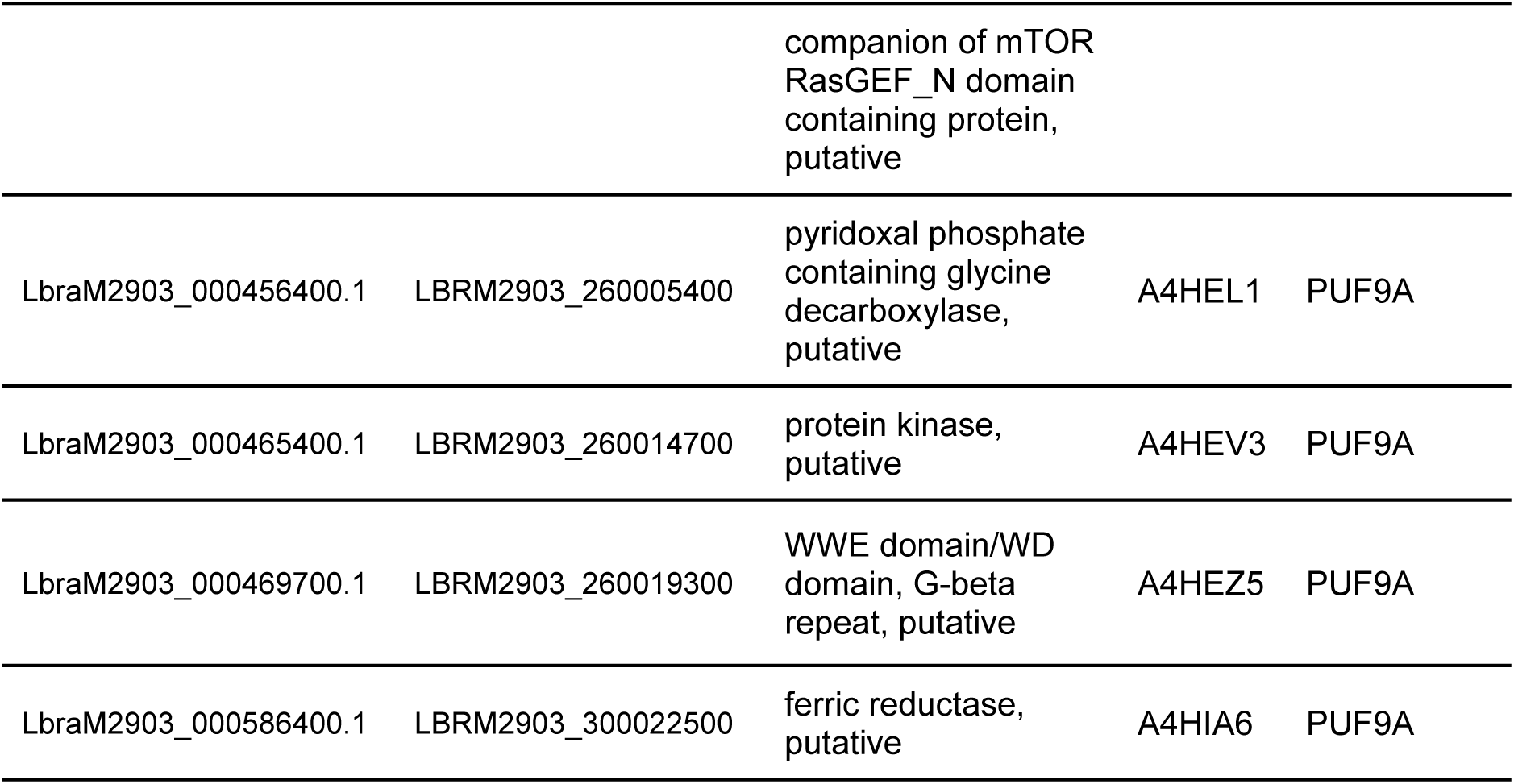
Differentially expressed genes co-expressed with mRBPs overexpressed in *L. braziliensis* resistant to SbIII, with physical interaction corroborated experimentally.

To explore direct regulatory potential, we scanned the 3’ UTRs of DEGs co-expressed with DRDB3 using its known *Trypanosoma brucei* experimentally validated binding motif (CTTTCT), as described by Ray et al. (2013) (38). Nine genes contained this motif in their 3’ UTRs, among them four encoding hypothetical proteins and other six annotated the following proteins: a P-type cation-transporting ATPase superfamily, subfamily IIA, SERCA-type; T-complex protein 1 subunit gamma; a zinc-containing-like protein; an ortholog of TP53I3; an Alba domain-containing protein; and an Anti-Müllerian hormone (Table 3).

**Table 3.** DEGs with 3’ UTR binding motif of DRBD3.

| Our annotation | TritypDB | Product | Uniprot |
| --- | --- | --- | --- |
| LbraM2903_000034500.1 | LBRM2903_040005000 | calcium-translocating P-type<br>ATPase | A4H3S2 |
| LbraM2903_000158600.1 | LBRM2903_130008100 | Alba, putative | A4H734 |
| LbraM2903_000170000.1 | LBRM2903_130020400 | hypothetical protein | A4H7E3 |
| LbraM2903_000259200.1 | LBRM2903_190008100 | hypothetical protein | A4H9V9 |
| LbraM2903_000381600.1 | LBRM2903_230005000 | hypothetical protein | A4HCL4 |
| LbraM2903_000392400.1 | LBRM2903_230018200 | alcohol dehydrogenase, zinc-containing-like protein | A4HCV8 |
| LbraM2903_000396500.1 | LBRM2903_230023100 | T-complex protein 1, gamma<br>subunit, TCP1g | A4HCZ3 |
| LbraM2903_000396600.1 | LBRM2903_230023200 | hypothetical protein | A4HCZ4 |
| mRNA_LbrM2903_26_233 | LBRM2903_260027300 | Anti-Mullerian hormone, N-terminal region containing<br>protein, putative | A4HF75 |

## DISCUSSION

Leishmaniasis, caused by protozoan parasites of the genus *Leishmania*, affects over a million people annually (39–41). While traditionally endemic to tropical countries, its geographical range is expanding with climate change. (42, 43). Current therapies are limited, often fail due to the emergence of resistant strains, and the development of new drugs remains scarce (4–6). In this context, low-cost computational approaches can generate insights to guide targeted experimental research, as demonstrated through the integration of updated predictive tools and databases (44).

One of the fundamental processes in all living organisms is the regulation of gene expression. Here, we present the most comprehensive comparative genomic and transcriptomic analysis of RNA-binding proteins (RBPs) and regulatory networks in *Leishmania* to date, revealing conserved regulatory elements, lineage-specific adaptations, and their involvement in stress responses and drug resistance. By re-annotating 33 genomes from 19 *Leishmania* species, including previously uncharacterized non-reference strains, we mapped the genomic repertoire of RBPs with unprecedented resolution. Our pan-genome analysis revealed a highly conserved core of RBPs, as well as subgenus-specific expansions. Importantly, we identified orthologs of RNA modification enzymes (e.g., NAT10, TRMT6/61A, NSUN2) conserved across all analysed species, despite the absence of canonical m6A writers. This divergence suggests that *Leishmania* may have evolved unique epitranscriptomic mechanisms, representing a fertile ground for future exploration.

Pan-genome analysis of protist genomes is challenging because there are few available genomes that capture their full diversity (45). Nonetheless, fungal pan-genome studies have shown that a particular species core genome may include about 80 to 90% of all protein-coding genes (46). This pattern was observed in our work when we analysed the pan-genomes of *L. major*, *L. braziliensis,* and *L. donovani* individually. However, this level of conservation decreases in species within the same subgenus, where gene conservation ranges from 20 to 31%. Moreover, when comparing the entire genus, only around 9% of the total genes are part of the core genome. Importantly, our comparative genomic analysis allowed us to characterise genomic features with unprecedented detail. Specifically, our study of putative regulatory elements revealed a high level of conservation in RBPs, which are essential for modulating mRNA translation and stability (22).

Similar to other trypanosomatids, *Leishmania* mainly regulates gene expression through post-transcriptional mechanisms (47–50), largely mediated by RBPs and already known to be linked to determining the cellular fate in *Leishmania* (47, 51). These proteins control multiple layers of regulation, including mRNA processing (such as capping, trans-splicing, or polyadenylation) (25), stabilising or destabilising mRNAs (25, 52), and adding and controlling mRNA modifications, such as N6-methyladenosine (m6A) or N4-acetylcytidine (ac4C) (35–37). Using a combination of homology searches and the hidden Markov model–based domain detection, we generated the most extensive genome-wide catalog of RBPs in *Leishmania* to date. The total numbers we identified were consistent with large-scale proteomic studies in *L. mexicana* (26) and exceeded tenfold those captured by poly(A)-bound RBP mass spectrometry (MS) datasets in *L. donovani* (*27*). However, a study by Kalesh et al. (2022) combined the OOPS method with TMT labelling-based quantitative proteomic MS in *L. mexicana* and identified more than 2,000 RBPs (47). Concerning RBP subtypes, and similar to previous search strategies, we detected non-coding RNA-binding proteins such as snoRNA binders (snoRBPs), which are required for 18S rRNA processing (53). snoRNAs have recently been linked to responses to metabolic and inflammatory stresses, as well as liver cancer (54, 55), implying that modulating or inhibiting snoRBP activity could be advantageous in treating these conditions. For example, the U3 small nucleolar RNA-associated protein 6 (UTP6) has been identified as a potential vaccineable virulence factor in *L. braziliensis* (56).

In trypanosomatids, mRBPs regulate translation through dynamic interactions with their target mRNAs (22, 57). We identified several genes encoding proteins that directly regulate translation or perform chemical modifications on mRNAs, such as methylation or acetylation. Key domains found in the putative mRBPs we identified include the Pumilio (PUF) domain, CCCH zinc finger (zf-C3H) domain, and RRM domains. We identified ten homologs of RBPs containing a PUF domain that are conserved across all *Leishmania* species. This finding is consistent with previous reports by Folgueira et al. (58); however, our revised annotation allowed us to identify PUF9 in *L. braziliensis*, a protein that was not previously detected in this genome. The RRM domain-containing proteins are related to mRNA stability and are primarily associated with the fine-tuning of translation, promoting adaptation and survival in response to environmental challenges (11). De Gaudenzi et al. estimated that the number of RRM domain-containing proteins in *L. major* is around 80 (59), a number corroborated in our work. Likewise, we identified very similar numbers of RRM domain-containing proteins in all the different *Leishmania* species. Importantly, two of these proteins, which are present in all *Leishmania* genomes, are putative homologs of proteins crucial for developmental control in *T. brucei*, RBP6 and RBP10 (60, 61). Regarding the proteins with a zf-C3H domain, these proteins are known to be essential in trypanosomatid development and differentiation (34, 62, 63). Notably, we identified putative orthologs of proteins such as ZFP2, ZC3H20, and ZC3H11 in all studied *Leishmania* genomes, which are essential components of the differentiation machinery in *Trypanosoma*.

mRNA nucleotide modification has an essential impact on gene expression (64). Twelve different chemical modifications have been identified in mRNA transcripts, which can occur in 5’ or 3’ untranslated regions (UTRs), coding regions, and introns of mRNAs (64). The most prevalent modification in eukaryotic cells is the N6-methyladenosine (m6A) (65). It has been documented that m6A is involved in the regulation of mRNA processing and expression, as well as splicing, translation, and stability (66, 67). In opisthokonta, m6A modifications are primarily performed by METTL3 and METTL14 (69, 70), and, alternatively, also by METTL16 (68). In trypanosomatids, the m6A modification has been described as enriched in the poly-A tails, a unique characteristic for these organisms. (36, 37). Despite this, orthologs to the m6A writers in trypanosomatids have not yet been identified (35, 69). The methodology employed by Maran et al. (2021) only utilised sequence homology to identify orthologs. However, although our approach proposes a more robust search by incorporating not only sequence homology but also domain searches using the Pfam database, it was still not possible to identify orthologs of METTL3, METTL14, or METTL16 in *Leishmania* species. This suggests that the ortholog of this methyltransferase is highly divergent compared to its counterparts in other eukaryotes (35), or this organism uses a noncanonical m6A writer enzyme.

Although m6A writer orthologs were not identified, orthologs associated with other modifications, such as N4-acetylcytidine (ac4C), were successfully located in all *Leishmania* species. The ac4C mRNA modification plays a crucial role in enhancing translation efficiency and is catalysed by the enzyme NAT10 (70, 71). Recent studies have demonstrated that NAT10 is essential for cell survival in various types of cancer. Its overexpression is linked to poor prognosis in patients with head and neck squamous cell carcinoma (72–74). In trypanosomatids, NAT10 orthologs were first described by Maran et al. (2021). They analysed the RNA helicase and N-acetyltransferase domains, identifying several mutations in the leishmanial NAT10 ortholog (35). In our study, we replicated this analysis by including species from different subgenera and causative agents of diverse clinical forms of leishmaniasis. Our findings indicate subgenus-specific substitutions in the nucleotide-binding site of the RNA helicase domain. Additionally, the pattern of substitutions in the Acetyl-CoA binding site of the N-acetyltransferase domain suggests a subgenus-specific adaptation for these motifs. A recent study by the Moretti group highlighted the essential role of NAT10 for the survival of *L. mexicana*, further implying it could be a promising therapeutic target for leishmaniasis (75). Remodelin, an inhibitor of NAT10, has been tested in various contexts as a chemotherapy agent that binds to the acetyl-CoA binding site of NAT10, inhibiting its acetyltransferase activity (76, 77). We compared the amino acids at the interaction site of Remodelin in human NAT10 (77) with their orthologs in *Leishmania*, noting that these amino acids are conserved. This suggests that Remodelin could potentially serve as an antileishmanial agent.

In addition to identifying potential regulators of gene expression, we also analysed their expression profile in *L. braziliensis* across various developmental stages and stress conditions caused by temperature shifts or exposure to antiparasitic agents (SbIII).

Several genes were overexpressed at each developmental stage. Notably, we observed a significant change in the expression of the ZC3H20 gene in the amastigote form of *L. braziliensis*. The product of this gene in *T. brucei* stabilises mRNAs encoding some procyclic-specific membrane proteins and is mainly expressed in procyclic forms (63, 78). However, the ZC3H20 gene is overexpressed in the amastigote form of *L. braziliensis*, suggesting that its product may have a different role compared to that of *T. Brucei.* Additionally, overexpression of RBP6 has been mainly associated with metacyclogenesis, as well as with the expression of genes coding for variant surface glycoprotein (VSG) and infectivity in *Trypanosoma* (62, 79–81). We observed RBP6 overexpression in metacyclic promastigotes relative to amastigotes of *L. braziliensis,* indicating that the gene product of this gene likely fulfils a similar function in metacyclogenesis in both *Leishmania* and *Trypanosoma*. Regarding ZC3H39, it is associated with mRNA stabilisation during the lifecycle, response to nutritional stress, and respiratome (23, 82). In *T. brucei,* the expression of the ZC3H39 gene remains constant throughout the developmental stages (19), a phenomenon also observed in *L. braziliensis*, with only a slight increase during the amastigote stage. PUF2 is a repressive mRBP in trypanosomes, necessary for normal growth of bloodstream forms (83). Our analysis revealed PUF2 overexpression in the amastigote stage of *L. braziliensis*, implying that its product is involved in this phase of development. PUF6 is a repressor that promotes mRNA degradation in *L. infantum* and *Trypanosoma*; however, it is non-essential for parasite growth (84). Dallagiovanna and colleagues (2008) reported PUF6 overexpression in the metacyclic form of *T. brucei* (85). We observed PUF6 overexpression in metacyclic promastigotes of *L. braziliensis*, indicating that its expression may be necessary for metacyclogenesis in trypanosomatids. Recently, Marucha and Clayton (2020) described PUF3 as a non-essential protein, although required for fine-tuning gene expression during the differentiation of the procyclic form of *T. brucei* (86). We also found PUF3 overexpressed in the procyclic form of *L. braziliensis*, which could suggest a functional relationship between these orthologs in trypanosomatids.

The interest in understanding the role of RBPs in drug resistance has grown, particularly in cancer and bacterial antibiotic resistance (90, 91). However, the role of RBPs in antileishmanial drug resistance still needs clarification. We found that the mRBPs DRBD3, ZFP2, and PUF9A are overexpressed in *L. braziliensis*-resistant promastigotes treated with SbIII. DRBD3 is a 3’ UTR binding mRBP which modulates the stress response, stabilising their target mRNAs in *T. brucei* (87–89). In *L. mexicana,* the expression of DRBD3 peaks in the metacyclic form and decays in amastigotes and procyclic forms (26). This phenomenon was also observed in *L. braziliensis*. However, its relationship with drug resistance has not been documented. We conducted a co-expression network analysis to explore the potential association of DRBD3, ZFP2, and PUF9A RBPs in drug resistance. This approach has previously been used to identify the role of RBPs as potential regulators and prognostic markers in myelopoiesis and leukaemia, as well as in lung squamous cell carcinoma, colorectal cancer, and lung adenocarcinoma (90–93). Our findings showed that these three RBPs are positively co-expressed with other 287 genes in promastigotes that are resistant to SbIII. Among these co-expressed genes, we identified an aminopeptidase, whose ortholog in *L. donovani* has been linked to resistance to SbV. (94). Importantly, aminopeptidases catalyse the removal of N-terminal amino acid residues from peptides and proteins and are proposed as emerging anti-parasite targets (95). Additionally, another gene that co-expresses with DRBD3, PUF9A, and ZFP2 is cytosolic leucyl aminopeptidase (LAP). LAP has mainly been used as a therapeutic target for treating malaria and toxoplasmosis (96, 97). In *T. brucei,* this enzyme is not essential for the survival of promastigotes in both in vivo and in vitro experiments. However, LAP does play a role in protein degradation during nutrient starvation (98). In *L. donovani,* overexpressing LAP was associated with a phenotype of resistance to α-difluoromethylornithine (DFMO) (99), suggesting that LAP stabilisation is vital not only for nutrient starvation but also for drug resistance in trypanosomatids. A recent study has also demonstrated that peptidomimetics can inhibit their enzymatic activity in *L. donovani,* opening up the possibility of developing new therapeutic strategies (100). The chaperonin TCP20 is also co-expressed with DRBD3, PUF9A, and ZFP2. In miltefosine-resistant *L. donovani,* TCP20 is downregulated (101, 102), while it is overexpressed in SbIII-resistant promastigotes of *L. braziliensis*. Another chaperone affected in SbIII-resistant *L. braziliensis* is DnaJ, which, similar to TCP20, is downregulated in miltefosine-resistant *L. donovani* (102). DnaJ is overexpressed in SbIII-resistant *L. infantum* (103), suggesting these two chaperones have opposite roles in resistance to both drugs. The pteridine reductase 1 (PTR1) also shows positive coexpression with three of our target RBPs. PTR1 plays a vital role in pteridine salvage and in antifolate resistance in *L. major* (104, 105). PTR1 is an NADPH-dependent biopterin reductase and is involved in oxidative stress defence within the macrophage (106, 107). In a proteomic analysis of antimony-resistant *L. braziliensis*, PTR1 was found to be overexpressed (108). Additionally, overexpression of PTR1 was observed in methotrexate-resistant *L. major* (106, 109).

Our protein-protein interaction analysis within the co-expression network of SbIII-resistant promastigotes has enhanced our understanding of potential proteins actively involved in the resistome, linked to the expression of DRBD3, PUF9A, and ZFP2. The 3-hydroxyacyl-ACP dehydratase (HTD2) is a protein participating in mitochondrial fatty acid synthesis in trypanosomatids, encoded by the LmjF07.0430 gene in *L. major* (110, 111). We observed that the orthologue of HTD2 in *L. braziliensis* is co-expressed with DRBD3, PUF9A, and ZFP2, and there is experimental evidence confirming its physical interaction with DRBD3. FabZ encodes HTD2 in *Plasmodium falciparum*, where small-molecule inhibitors of FabZ have been successfully used against this parasite, resulting in the inhibition of fatty acid synthesis and growth *in vitro* (112). However, the potential of HTD2 as a therapeutic target in trypanosomatids remains to be investigated. Similarly, we identified interactions between SEC61α and DRBD3 and PUF9A in SbIII-resistant *L. braziliensis.* SEC61 is a protein complex required for secretory protein translocation and integral membrane protein insertion in *S. cerevisiae* (113). It is composed of three subunits: SEC61α, SEC61β, and SEC61γ. Recently, two natural products were studied as inhibitors of SEC61α, showing promise as potent agents for treating diseases such as cancer (114, 115). In *T. brucei,* silencing SEC61 induces spliced leader RNA silencing and leads to programmed cell death (116), suggesting it could also be a promising therapeutic target against trypanosomatids. S-adenosylmethionine synthetase (METK) is another gene overexpressed in promastigotes of *L. braziliensis* SbIII-resistant, and it is co-expressed with PUF9A, DRBD3, and ZFP2. METK plays a role in polyamine synthesis and is involved in the synthesis of glutathione and trypanothione in trypanosomatids (117, 118). In a proteomic study of *L. panamensis*, METK levels were found to be overexpressed in clinical isolates resistant to SbIII (119), suggesting a stabilising role of these RBPs on METK mRNA. Additionally, a protein interaction between METK and PUF9A was reported in the databases. Bhattacharya et al. (2019) discovered that Sinefungin, an analogue of S-Adenosylmethionine, mainly targets the METK protein, thereby inhibiting the growth of *L. infantum* (120). This indicates that such an analogue substrate could potentially be used in species resistant to other treatments, such as antimony. We also found that RNA Binding Protein 5, encoded by RBP5, interacts with DRBD3. RBP5 in *T. brucei* is crucial for cell proliferation and survival, as well as regulating intracellular iron levels (121–123). Recently, RBP5 has been shown to play a regulatory role during *Leishmania* mating (124). However, our evidence provides the first support for its potential role in antimony resistance.

Finally, we mapped the binding motif of DRBD3 with all the 3’ UTRs of their co-expressed genes, aiming to identify a post-transcriptional gene regulatory network. Our results revealed the identification of 10 genes whose mRNAs are very likely to be stabilised by DRBD3 in *L. braziliensis* promastigotes resistant to antimony, suggesting that these genes could play an essential role in the mechanism of resistance to this drug. This is supported by the gamma subunit of the T-complex protein 1 (TCP1-gamma) and Alba protein 1 (ALBA1), genes that are constitutively expressed in *L. infantum* (125) and overexpressed in SbIII-resistant *L. amazonensis* (126). This finding suggests that DRBD3 could play a crucial role in SbIII resistance in *Leishmania*, making it a promising marker for antimony resistance and a potential target for future therapeutic studies.

In summary, this work provides the most comprehensive comparative analysis of RNA-binding proteins and RNA modification machinery across Leishmania to date, integrating genomic annotation, expression profiling, and network analysis. We demonstrate that RBPs are highly conserved yet exhibit context-specific expression patterns linked to development, stress responses, and drug resistance. Importantly, we reveal that key mRBPs, including DRBD3, PUF9A, and ZFP2, are associated with antimony resistance and co-expression networks enriched for potential therapeutic targets such as aminopeptidases, NAT10, PTR1, SEC61, HTD2, and METK. These findings emphasize the central role of post-transcriptional regulation in parasite adaptation and point to RBPs as both biomarkers and candidates for drug development. Future work should focus on experimental validation of the predicted regulatory interactions, binding motifs, and protein–protein associations described here, using approaches such as CLIP-seq, RIP-seq, and functional knockouts. Beyond validating individual targets, integrating RBP studies with chemical biology and high-throughput drug screening will be critical to translate these discoveries into therapeutic opportunities. By highlighting RBPs as master regulators of gene expression and resistance, our study provides both a framework for future experimental exploration and a roadmap for the rational design of new antileishmanial strategies.

## METHODS

### Datasets acquisition

Genome sequences of *Leishmania* strains were obtained from TriTrypDB (127) release 66 in January 2024. Raw RNA sequencing data were retrieved from the NCBI Sequence Read Archive (SRA) (128) in December 2023, encompassing three publicly available studies. The first dataset corresponded to *L. braziliensis* M2903 (MHOM/BR/75/M2903), generated by Ruy et al. (2019) (129), with accession number PRJNA494068. This study included samples from axenic amastigotes, procyclic, and metacyclic promastigotes at various developmental stages. The second dataset (accession PRJEB31092) was published by Patino et al. (2019) (130) and contained *L. braziliensis* promastigotes that were resistant or susceptible to trivalent antimony (SbIII). The third dataset, reported by Ballesteros et al. (2020)(131), comprised temperature-stressed *L. braziliensis* promastigotes (accession PRJEB31852).

### Identification of RNA-binding proteins

To determine the repertoire of RBPs in each *Leishmania* genome, we used 791 Hidden Markov Models (HMMs) of RNA-binding domains (RBDs) obtained from EuRBPDB (132). First, we mapped all predicted proteins against these HMMs with hmmscan using an E-value threshold of 1E-04 to assign membership to a specific RBP family. In parallel, proteins were searched against Pfam (133) to confirm the presence of RDB-associated HMMs.

We further performed a homology-based search to identify mRPBs, with a focus on those involved in mRNA modification, stabilization, or repression. A reference set of known trypanosomatid mRBPs, previously described in the literature (25, 27, 28, 35, 134, 135), were downloaded from eggNOG 5.0 (136) and compared against the predicted proteomes of *Leishmania* genomes using DIAMOND (137), applying thresholds of 1e-05 for E-value, 70% query coverage, and 50% sequence identity. Multiple sequence alignments of candidate hits were generated using Clustal Omega v1.2.4 (138, 139) and visualised using Jalview v2.11.2.2 (140).

To define putative binding sites for an mRBP, we used a homology-transfer approach. mRBP binding motifs experimentally characterized in trypanosomatids (38) were compared with *Leishmania* orthologs, using 75% sequence identity and query coverage cutoff to infer motif conservation. Once binding motifs were assigned across the 32 genomes, we scanned entire genomes using FIMO (141) from the MEME suite v5.4.1 (141, 142). An mRBP was considered to regulate a gene if its binding motif was found within the last 750 nucleotides of the gene’s 3’UTR (143, 144).

### Comparative analysis of RBPs in Leishmania genomes

To perform a comparative genomic analysis, we employed a bidirectional best hit algorithm for orthologs clustering using ProteinOrtho v5.11 (145) with default parameters for 32 *Leishmania* spp predicted RBPs analyzed. Functional analysis of core and pan-RBPome was performed using ClusterProfiler v4.4.3 (146).

### RNA-seq data processing and differential expression analysis

Raw RNA-seq data were processed following a previously established protocol (147). In summary, FastQC v0.11.8 (https://www.bioinformatics.babraham.ac.uk/projects/fastqc/) and Trimmomatic v0.36 (148) were utilized to assess and eliminate low-quality reads. Subsequently, the remaining reads were aligned to the target genomes using Bowtie2 v2.3.4.1 (149). Count matrices for each experiment were generated using featureCounts v2.0.6 (150). Normalised counts in TMM were then computed using EdgeR v4.0.3 (151). Batch effect removal was achieved through a parametric empirical Bayes framework adjustment with SVA v3.50.0 (152). Differential gene expression analysis was conducted using DESeq2 v1.32.0 (153), with a significance threshold of false discovery rate (FDR) or adjusted p-value < 0.001, and ∣log2FC∣ > 0.75. Additionally, GO enrichment analysis was performed using the Cytoscape app BiNGO v3.0.5 (154) in Cytoscape v3.10.1 (https://cytoscape.org/).

### Construction of context-specific co-expression networks analysis

To determine context-specific co-expression patterns, we adapted the approach by Kuijjer et al. (2019) (155). Only genes with expression >0.5 counts per million (CPM) in at least 90% of the libraries were considered. Pearson correlations were computed for all pairs of expressed genes across the combined dataset. Available RNA-seq data for *Leishmania* parasites corresponded to *L. braziliensis* samples from all developmental stages, including amastigote, procyclic promastigotes, and metacyclic promastigotes (129). In addition, we included *L. braziliensis* promastigotes samples grown at 24°C, 26°C, 28°C, and 30°C (131). Furthermore, to assess antiparasitic resistance, we also analysed resistant and susceptible *L. braziliensis* samples to trivalent antimony (SbIII) (130). Gene pairs correlations were computed across the entire dataset and classified as total correlations (RTotal). Then, we measured the contribution to the R_Total_ of a specific context by calculating the Pearson correlation for each pair of genes using all samples except those from the context of interest (R_Context_) and subtracting this correlation from R_Total_. For example, in the context of drug resistance, RContext is calculated for each gene pair using all samples except those from drug-resistance experiments. The contextualised weight or perturbation (P) between each pair of genes is then calculated as P = R_Total_ - R_Context_. After that, we adjusted perturbation values as follows: P_Rescaled_ = 1 ― (2((*x* ― *min*)/(*max* ― *min*))), where x is the perturbation value, min is the minimum perturbation value, and max is equal to the maximum perturbed value for each context studied. Next, the perturbation matrix was clustered using a hierarchical clustering algorithm, and the resulting dendrogram was cut to determine the optimal number of modules using the Dynamic Cut Tree package (152), yielding the set of strongest correlations between pairs of genes.

To validate our method, we evaluated the expression patterns of a set of gene markers of the amastigote developmental stage based on the hypothesis that genes with similar expression patterns should be co-expressed and part of the same modules. We built a list of amastigote markers based on the literature (156–158), 202 protein sequences that were aligned against *L. braziliensis* proteins with BLAST, and annotated two proteins as homologs if they share at least 50% sequence identity, 50% query coverage, 50% target coverage, and their alignment has an e-value of at most 1e-5.

To determine the occurrence of each gene pair across different contexts, we constructed a module co-occurrence network. In this network, we compare each gene pair to decide whether or not they are co-expressed across all networks within each context. For instance, in a context *C* with *x* = 3, where *x* represents the number of networks comprising this context, the co-occurrence score for the gene pair A-B is evaluated. The maximum possible score in this example is 3, indicating that both genes co-occur in the same module across all three networks in that context. Conversely, a score of 0 signifies that the genes do not co-occur in any module within this context and are not co-expressed.

### Protein-protein interaction analysis

Protein-protein interactions (PPIs) were evaluated using the STRING database v11.5 (https://string-db.org/) (159). Co-expressed gene sets identified from our networks were queried to determine whether their encoded proteins exhibited experimentally supported interactions. An initial combined confidence score threshold of 0.1 was applied, followed by manual evaluation of evidence for direct physical interactions.

## ACKNOWLEDGMENTS

Powered@NLHPC: This research was partially supported by the supercomputing infrastructure of the NLHPC (CCSS210001).

---

**Supplementary Figure 1.** Expression patterns of selected mRBPs in promastigotes of Leishmania braziliensis under temperature stress. Here, we identified three groups. In the first group, we observed a peak in the expression of mRBPs genes at 26°C (pro26). Contrary, in group 2, we observed that at 26°C, mRBPs have less expression. Finally, group 3, composed of NAT10 and HNRNPH, presents their expression peak at 24°C (pro24).

